# Reactive oxygen species alter ex vivo dopaminergic and purinergic signaling and microglia physiology in the nucleus accumbens

**DOI:** 10.64898/2026.05.01.722235

**Authors:** Hillary A. Wadsworth, Justin A. Webb, Erin B. Taylor, Eliza R. White, Lauren H. Ford, Lydia R. Hawley, Brayden J. Parker, Stephen T. Jones, Sara C. Linderman, Christopher J. Galbraith, Derek D. Langford, Cody A. Siciliano, Jason M. Hansen, Scott C. Steffensen, Jordan T. Yorgason

## Abstract

Microglia are the brain’s resident immune cells and rapidly migrate toward sites of neuronal stress and neurotransmitter imbalance. The molecular cues directing microglia movement remain poorly understood, particularly in brain regions involved in reward processing, including the nucleus accumbens (NAc). Dopamine (DA) dysregulation and production of reactive oxygen species (ROS) are hallmarks of substance use disorders and neurodegenerative diseases. Dopamine and adenosine triphosphate (ATP) are co-released in the NAc, but the effects of ROS on ATP/DA co-transmission and microglia surveillance remain unknown. Using multiphoton imaging in brain slices with GFP-labeled microglia, we found that ATP consistently and non-selectively drives microglia chemoattraction, whereas DA chemoattraction only occurs in a subset of microglia, likely through D1 receptor activation. Microglia become more reactive via LPS application, and in associated experiments, LPS results in enhanced DA/ATP release measured via fast scan cyclic voltammetry. ROS production also transitions microglia to a more reactive state, but inhibits DA release, with mixed effects on ATP release. These findings identify ATP- and DA-mediated mechanisms for NAc microglia chemoattraction and opposing immune dysregulation (via LPS or ROS) of DA terminal function.

## Introduction

The nucleus accumbens (NAc) of the mesolimbic dopamine (DA) system is responsible for learned associations for motivation and reward (Salgado & Kaplitt, 2015) and receives dopaminergic projections from the ventral tegmental area (VTA) (Ikemoto, 2007). VTA DA neurons express the vesicular nucleoside transporter (VNUT), which co-packages ATP with DA into vesicles (Ho et al., 2015) to co-release ATP with DA from VTA projections (Borgus et al., 2021; Linderman et al., 2026). The effects of DA on striatal neuronal circuits are well characterized, though little is understood about how DA terminals influence local microglia cell function and, conversely, how microglia influence DA terminals.

Microglia are the brain’s resident immune cells. They exhibit a ramified morphology, where branch-like processes continuously extend and retract to survey nearby tissue (Goldmann et al., 2016), particularly in response to purinergic signaling (Wollmer et al., 2001). This surveying behavior is a form of “resting” activity, and microglia from cultured or dissociated preparations lose process surveillant behavior while maintaining motility (Boche et al., 2013; Gipson et al., 2021). Microglia express DA and purine receptors (Färber et al., 2005; Illes et al., 2020; Schwarz et al., 2013), though expression varies by brain region (Borst et al., 2021; Keren-Shaul et al., 2017). Microglia ramification is associated with non-inflammatory states and may reflect local activity from DA terminals. On the other hand, microglia can change rapidly to a more enlarged amoeboid form in response to injury or inflammation. This morphological shift reflects a change in function and has been shown to be associated with increased release of various pro-inflammatory cytokines (Qin et al., 2005). Microglia processes are also highly reactive to local activity, particularly local release of nucleosides (Abiega et al., 2016; Orr et al., 2009). P2X and P2Y purinergic receptors expressed on microglia induce microglia morphology changes upon activation (Colella et al., 2018; Illes et al., 2020), with ATP acting as a chemoattractant for processes (Honda et al., 2001; Nasu-Tada et al., 2005) and uridine 5’-diphosphate causing microglia to become phagocytic (Koizumi et al., 2007). While ATP-sensing by microglia is typically described in the context of cellular damage as a damage-associated molecular pattern (DAMP) (Abbracchio et al., 2009), the activity and reactivity of microglia in non-inflammatory states, and the possibility of co-release of DA and ATP, suggests there is an unexplored effect of co-transmitter effects on microglial processes and motility. Therefore, the current study examines the effects of DA and ATP on NAc microglial chemotactic motility.

Microglia are also major generators, detectors, and regulators of reactive oxygen species (ROS) signaling. Microglia express NADPH oxidase (NOX), which produces ROS during inflammation through NOX2 and NOX4 activation (Simpson & Oliver, 2020; Zhang et al., 2014; Zhang et al., 2016). To sense ROS, microglia express the H_2_O_2_-sensitive nonselective cation channel transient receptor potential melastatin 2 (TRPM2) (Jeong et al., 2017; Malko et al., 2019; Miller & Zhang, 2011), though there may be cell heterogeneity in receptor expression. Microglia also express various peroxidases that can rapidly terminate ROS signaling, including glutathione peroxidases (GPX) and selenoproteins (Hirrlinger et al., 2000; Meng et al., 2019), which may be a mechanism for fine-tuning the localization of ROS signaling. Through these various mechanisms, cells can produce, sense, and rapidly terminate ROS signals, though the extent of NAc microglia involvement in these mechanisms is unknown. Dopamine release is inhibited by ROS such as H_2_O_2_, an effect thought to be through direct interactions with KATP channels on DA terminals (Patel & Rice, 2012). However, previous work has not considered the site of pharmacological effects, nor the presence of K_ATP_ channels on microglia (McLarnon et al., 2001; Ortega et al., 2012). Prior work in our lab has demonstrated a role for ROS production in the DA-enhancing effects of methamphetamine, cocaine, and nicotine (Hedges et al., 2018; Jang et al., 2015; Jeon et al., 2025; Torres et al., 2021), but the effects of local ROS production on DA and ATP corelease and subsequent effects on NAc microglia have remained unexplored. Therefore, the present work also examines the effects of ROS production on NAc neuroimmune circuitry.

## Methods

### Animal Care

Male and female adolescent (PND 30-69, n=34) young adult (PND 70-180, n=62), and mature adult (PND 181-225, n=2) C57BL/6J background (WT and transgenic) mice were used throughout. No animals were excluded based on age and there were no obvious age dependent effects. Animals had *ad libitum* access to food and water and maintained on a 12:12 hour light/dark cycle. Csf1r-EGFP expressing macrophage FAS-induced apoptosis (MaFIA) mice (RRID:IMSR_JAX:005070) were used for all imaging experiments. In the central nervous system (CNS), Csf1r is primarily expressed in microglia (Erblich et al., 2011). In MaFIA mice, IBA1 has been reported to co-label in all Csf1r-EFGP cells in the brains of control mice (Gabrusiewicz et al., 2015). However, Csf1r expression has also been reported in limited amounts in CNS-associated macrophages, certain neural progenitor cells, and neuronal subpopulations (Hu et al., 2021). MaFIA mice were presently used for their Csf1r-EGFP labeling of macrophage-lineage cells, confirmed previously to be primarily IBA1+ microglia (Gabrusiewicz et al., 2015). As such, no additional validation was used herein to identify cells as microglia specifically. All protocols and animal care procedures were in accordance with the National Institutes of Health Guide for the care and use of laboratory animals and approved by the Brigham Young University Animal Care and Use Committee.

### Brain Slice Preparation

Mice were anesthetized with isoflurane (CHEBI:6015; Patterson Veterinary) before being decapitated with a razor blade and their brains carefully placed in pre-oxygenated (95% O2/5% CO2) artificial cerebral spinal fluid (ACSF; 35°C, pH ∼7.4). The ACSF consisted of (in mM): 126 NaCl, 2.5 KCl, 1.2 NaH2PO4, 1.2 MgCl2, 21.4 NaHCO3, and 11 D-glucose. The ACSF used for slicing was treated with ketamine (2 mM; NMDA receptor antagonist) to prevent excitotoxity. Coronal brain slices (containing NAc) were obtained with a vibratome (Leica VT1000 S; thickness of 220 µm).

### Drug Preparation and Administration

The following concentrations of drugs were used for the various experiments: ATP iontophoresis (10 mM; Sigma-Aldrich, St. Louis, Missouri), DA iontophoresis (1 mM; Cayman Chemical Company, Ann Arbor, Michigan), DA perfusion (100 µM; Cayman Chemical Company, Ann Arbor, Michigan), SCH23390 (1 µM; Cayman Chemical Company, Ann Arbor, Michigan), Sulpiride (600 nM; abcam, Cambridge, Massachusetts), glucose oxidase (1, 3, 10, and 30 mU/mL; Sigma-Aldrich, St. Louis, Missouri), and lipopolysaccharide from *Escherichia coli* O111:B4 (LPS; 1 µg/mL; Sigma-Aldrich, St. Louis, Missouri).

### Fast-Scan Cyclic Voltammetry (FSCV)

Dopamine and ATP release were detected concurrently using previously established FSCV protocols (Borgus et al., 2021; Yorgason et al., 2011). Briefly, a stimulating borosilicate glass electrode (P-87 Horizontal pipette puller; Sutter Instruments) was filled (1 mM KCl) and placed in the region of interest with electronic micromanipulators (Siskiyou). The recording electrode is a carbon fiber electrode (CFE) made by aspirating a carbon fiber (diameter ∼7 μm, Thornel T-650, Cytec) into a borosilicate glass capillary tube (TW150, World Precision Instruments), before heating and pulling on a horizontal puller to form a tight seal of glass around the carbon fiber. The carbon fiber is then trimmed (∼100-200 μm), filled with KCl (1 mM), and placed ∼150 μm away from the stimulating electrode. A voltage ramp is applied to the CFE (−0.4 to +1.5 V) at a scan rate of 400 V/s and a frequency of 10 Hz to measure DA and ATP corelease, with all measurements being made against a Ag/AgCl reference electrode (Borgus et al., 2021). Demon Voltammetry and Analysis software (Yorgason et al., 2011) was used to automate acquisition and facilitate analysis. Baseline experiments measure neurotransmitter release across a period of 20-40 minutes to establish stable baselines. For glucose oxidase experiments, a stimulation protocol of 10 Hz, 30 pulses, occurred 2 minutes apart with data acquired continuously (Patel & Rice, 2012). For 4-hour LPS experiments, stimulation was one pulse every 5 minutes.

### Multiphoton Imaging of Microglia in Brain Slices

A Ti-Sapphire Chameleon Discovery NX Laser (Coherent) is tuned to 920 nm (optimized for EGFP) with the power set to 3-5 mW (5-15% intensity with gain at 10%) and visualized using a 40x (0.8NA) water immersion objective (Olympus), as reported previously (Wadsworth et al., 2024). For experiments measuring microglia movement, collections were made every ∼2 minutes. The z step was set to ∼1 μm. For exogenously applied ATP/DA, iontophoresis was used while imaging microglia. ATP (10 mM) and DA (1 mM) were each placed in a glass electrode (1-2 μm, 3 MOhm) and a retention current was applied (ATP: +19pA, DA: −19pA) to prevent leakage. The electrode was then lowered down into the slice. The first 30 μm of brain slice surface were avoided for experiments to prevent issues associated with microglia morphology in this section of tissue (Etienne et al., 2019). Before iontophoresis, a stable baseline was recorded for 10-20 minutes to ensure no drift or microglial movement. ATP or DA was then applied once/minute for 10 minutes via iontophoresis pulses (iontophoretic potential; ATP: −200 nA, DA: +200 nA). The microglia were recorded for at least 60 minutes following iontophoretic application. To analyze microglial motility and morphology, the programs FIJI (Schindelin et al., 2012) and ICY (de Chaumont et al., 2012) were used following protocols from (Young & Morrison, 2018) and (Etienne et al., 2019). Briefly, for motility experiments FIJI was used to prepare files for analysis while ICY was used to measure fluorescence values in the area surrounding the iontophoretic electrode. For morphology experiments FIJI was used for processing and analyzing microglia before and after glucose oxidase and LPS treatment following the protocol from Young and Morrison (2018). Experiments were blinded during analysis to reduce experimenter bias. Images were filtered using FIJI’s Brightness/Contrast, Unsharp Mask, and Despeckle functions. Post-filtered images were thresholded and microglia manually selected by analyzers. After microglia were identified, the images went through a final cleaning process through FIJI’s Despeckle, Close, and Remove Outliers functions. FIJI’s Skeletal Analysis was used to automatically collect number of branches and branch length of the selected microglia. Diagrams showing experimental conditions were created with Biorender software (https://biorender.com/). Diagram of glucose oxidase H_2_O_2_ production made using Molview v2.4 open source software (MolView).

### Immunohistochemistry (IHC) and Confocal Multiplex Imaging

All IHC solutions are in phosphate buffered saline (PBS) solution. Mice were anesthetized with isoflurane (CHEBI:6015; Patterson Veterinary) and perfused (transcardial) with PBS followed by 4% paraformaldehyde (PFA). After dissection, brains were postfixed overnight in PFA at 4°C. Brains were then transferred to sucrose (30%) for an additional 48 h at 4°C for cryoprotection. Coronal sections (40 μm) were obtained from bregma +1.10 to +0.14 mm (AO 860 Sliding Microtome; American Optical) and left free-floating in PBS for labeling. A protocol from (Radtke et al., 2020) was adapted to fit the parameters of our experiment. Briefly, sections were washed twice for 10 minutes with PBS containing 1% animal serum and 0.4% Triton X-100 in PBS (PBS-T). Sections then were washed for 30 minutes with 5% animal serum and 1% Fc block in PBS-T. Primary antibodies were applied in 1% animal serum and 1% Fc block with 0.1% Triton X-100 in PBS, and sections were incubated overnight on an agitator at 4°C. Sections were then washed twice in 1% animal serum in PBS-T for 10 minutes before being mounted onto glass slides (Globe Scientific) and a coverslip attached with either Vectashield HardSet mounting medium (Vector Laboratories) or EverBrite True Black Hardset Mounting Media. Antibody concentrations were selected based on recommendations from suppliers. All IHC experiments used MaFIA mice, which express EGFP in all monocytes, including monocyte derived microglia (Burnett et al., 2006). Anti-GFP antibodies (Aves Labs, Cat. No. GFP-1020; 1:2000) were used to label microglia for imaging. Additional antibodies used include P_2_X_4_ receptors, (Invitrogen, pa5-77663; 10 µg/mL; CF405S), P_2_Y_6_ receptors (abcam; Cat. # ab92504; 1:200; CF405S), D1 receptors (Novus biotechne; Cat. # NB110-60017AF647; 1:1000; pre-conjugated to AF647), D2 receptors (Bioss; Cat. # bs-20729R-A647; 1:200, pre-conjugated to AF647), and TRPM2 (Alomone Labs, Cat. No. ACC-043; 1:100; CF647) and were conjugated to their respective fluorophores (Biotium for CF dye antibody labeling kits). Z-stack images of the NAc core were collected using Olympus IX81 laser-scanning confocal microscopes with the FV1000 and FV3000 Fluoview systems (Olympus) with a UApo /340 20×/0.75 objective.

*Detection of H_2_O_2_ via Amplex Red.* Amplex Red experiments were conducted in ACSF to measure glucose-oxidase-dependent H_2_O_2_ production. Experiments followed protocols from Invitrogen Amplex Red Hydrogen Peroxide/Peroxidase Assay Kit (Cat. No. A22188, Invitrogen, Carlsbad, CA). Briefly, an Amplex Red reagent stock solution, reaction buffer, horseradish peroxidase stock solution, and H_2_O_2_ calibration solution were prepared. Samples with chosen glucose oxidase concentrations in ACSF were compared to prepared samples with known H_2_O_2_ concentrations. Fluorescence was then measured and compared using a Spectramax iD3 (Molecular Devices) to determine the amount of H_2_O_2_ created by the varying concentrations of glucose oxidase.

### Statistical Analysis

DA- and ATP-release FSCV experiments were analyzed in Demon Voltammetry suite (Yorgason et al., 2011). Release was measured at peak oxidation currents. Clearance was measured as the time constant (tau) from a single exponential fit between the peak current and its return to baseline. To compare across multiple animals and slices, DA- and ATP signals were averaged (within subject) across the last three ACSF recordings to establish a baseline value that subsequent DA and ATP signals were normalized to. Thus, post-drug conditions were compared to pre-drug baseline conditions in the same slice for a within-subject experimental design. To ensure scientific rigor, analyzers were blinded for microscopy and electrochemistry analysis, and statistics, sample sizes, and replication were specifically designed to verify results. No experiments were excluded in the analyses, and the statistical unit (e.g. animal/slice/cell) is noted in the results for each comparison. In instances where more than one unit came from the same animal, this is described in the text. Statistical analysis was performed using Prism 5 (GraphPad). Significance for all tests was set at p < 0.05. Values are expressed as mean ± SEM. Significance levels are indicated on graphs with asterisks *, **, and *** corresponding to significance levels p < 0.05, 0.01, and 0.001, respectively.

## Results

### NAc microglia express D1 and D2 receptors and show chemoattraction for ATP and DA

IHC was used with NAc brain slices to establish the DA and ATP receptor expression profile in MaFIA mice, which express Csf1r-EGFP in monocyte derived macrophages. Cells labeled with EGFP in the NAc were assumed to be microglia, as prior work with this mouse identified IBA1+ labeling in all Csf1r-EGFP cells under control conditions (Gabrusiewicz et al., 2015) and CNS-associated macrophages are most commonly found in CNS interfaces including the meninges, perivascular space, and choroid plexus (Prinz et al., 2021). The role for purinergic receptors in microglia motility outside the NAc is well established (Haynes et al., 2006; Honda et al., 2001), and we observe purinergic P_2_X_4_ (**Supplementary Fig. 1A**) (Wadsworth et al., 2024) and weaker expression of the uridine diphosphate (UDP) sensitive P_2_Y_6_ receptors (**Supplementary Fig. 1B**) on NAc microglia from fixed MaFIA brain slices. The expression and role of DA receptors in NAc microglia motility is less known, though DA receptor expression on leukocytes, monocytes, and microglia is highly variable and likely varies from cell to cell (Hui et al., 2025; Mather et al., 2008; Nelson et al., 2026). We observe DA D1 and D2 receptors throughout the NAc, with variable co-labeling on GFP+ microglia (**Fig. 1A**; CSF1R-GFP+ cells) in MaFIA mice, with strong labeling in non-GFP-labeled cells, which are putative medium spiny neurons (MSNs). In contrast to local expression on nearby cells (e.g., puncta on putative D1 MSNs), the D1R labeling on microglia appears more diffuse. Note that D2R labeling on microglia is more punctate and contrasts the diffuse labeling on putative D2 MSNs. Together these data indicate expression of DA receptors on NAc microglia.

**Figure 1:**
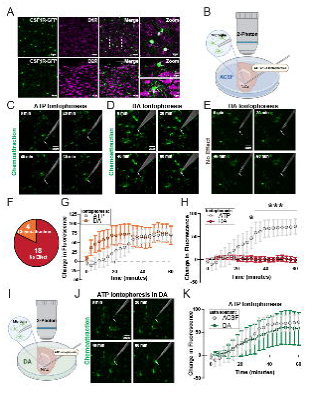
ATP and DA microglia chemotactic motility. **(A)** Immunohistochemical panel showing that D1 and D2 receptors are located on NAc microglia. **(B)** Schematic of experimental conditions for **C–H**. **(C)** Example experiment of ATP (10 mM) iontophoresis at time points 0, 20, 40, and 60 minutes. In this and all subsequent example iontophoresis experiments, the arrow represents the electrode tip, and a mask of the infrared channel is overlaid to show the approximate location of the iontophoresis electrode. **(D)** Example experiment of DA (1 mM) iontophoresis showing chemoattraction at time points 0, 20, 40, and 60 minutes. **(E)** Example experiment of DA (1 mM) iontophoresis showing no effect at time points 0, 20, 40, and 60 minutes. **(F)** 18/22 DA iontophoresis experiments did not show chemoattraction while 4/22 experiments did. **(G)** Group data showing the change in fluorescence at the electrode tip over time in cases where DA iontophoresis induced chemoattraction, with ATP data shown as a positive control for comparison. In experiments where DA chemoattraction was observed, microglia respond quicker to DA iontophoresis than to ATP. **(H)** Group data showing change in fluorescence over time for DA experiments, with ATP data shown for comparison. **(I)** Schematic of experimental conditions for **J–K**. **(J)** Example experiment of ATP iontophoresis with DA (100 µM) bath solution at time points 0, 20, 40, and 60 minutes. **(K)** Group data showing that perfusion of DA has no effect on microglia chemoattraction to ATP iontophoresis. Asterisks *,*** indicate significance levels p < 0.05 and p < 0.001, respectively. Group data are presented as mean ± SEM.

Next, GFP+ NAc microglia were visualized in live brain slices from MaFIA mice to determine chemoattractant motility (**Fig. 1B-I**) toward locally applied ATP (10 mM) or DA (1 mM) using iontophoresis. A schematic of the experiment is depicted in **Fig. 1B**. Baseline microglia positioning was determined before iontophoresis, and a change in fluorescence at the tip of the iontophoretic electrode was used to measure changes in microglia positioning near the chemical source. Robust microglia chemoattraction to ATP was consistently observed (**Fig. 1C, Supplementary Video 1**). Interestingly, DA iontophoresis had mixed effects (Chemoattraction example: **Fig. 1D, Supplementary Video 2;** No chemoattraction example: **Fig. 1E, Supplementary Video 3**), with chemoattraction in ∼22% of experiments (**Fig. 1F**), demonstrating that DA does not cause consistent chemoattractant of microglia compared to ATP. Interestingly, chemoattraction rates between ATP experiments and experiments where DA chemoattraction occurred were compared, and significantly higher rates of chemoattraction were observed to DA over ATP (**Fig. 1G**; two-way ANOVA; Iontophoretic solution, *F*_(1, 189)_ = 11.38, *p* = 0.0009; time, *F*_(20, 189)_ = 1.958, *p* = 0.0109; interaction, *F*_(20, 189)_ = 0.5142, *p* = 0.9585; ATP animals/slices n=6; DA animals/slices n=5). Unsurprisingly, when ATP experiments were compared to DA experiments, there were significant effects and interaction between experiments (**Fig. 1H**; Two-way ANOVA; Iontophoretic solution, *F*_(1, 540)_ = 160.2, *p* < 0.0001; time, *F*_(20, 540)_ = 4.488, *p* < 0.0001; interaction, *F*_(20, 540)_ = 5.237, *p* < 0.0001; ATP animals/slices n=6; DA animals n=14, slices n=22). Thus, ATP more consistently induces microglia chemoattraction, with DA causing chemoattraction for microglia in ∼22% of experiments.

We have established that DA chemoattraction occurs in a subset of microglia. However, it is possible that DA receptors on microglia could influence ongoing ATP chemoattraction. Therefore, the effects of bath-applied DA on iontophoretic ATP chemoattraction were examined. ATP iontophoresis experiments were conducted with simultaneous bath application of DA+ACSF (100 µM; **Fig. 1I, Supplementary Video 4**) or ACSF alone. Application of DA had no apparent effect on microglia ATP chemoattraction, though variability appeared to increase somewhat (**Fig. 1J-K**; two-way ANOVA; Perfusion solution, *F*_(1, 251)_ = 0.4661, *p* = 0.4954; time, *F*_(20, 251)_ = 1.974, *p* = 0.009; interaction, *F*_(20, 251)_ = 0.0606, *p* = 1.0; ACSF animals/slices n=6; DA animals n=7, slices n=8). Therefore, NAc microglia express DA D1 and D2 receptors, but DA chemoattractant effects are restricted to a subset of microglia and effects of DA on ATP chemoattraction were not observed, suggesting that NAc microglia prioritize ATP signaling.

### DA chemoattraction is absent during D1 but not D2 receptor antagonist application

Since DA chemoattraction is less frequent, but quite robust when observed, we next sought to determine if a specific DA receptor subtype was involved in DA chemoattraction. Therefore, DA iontophoresis was completed with concurrent antagonism of either D1 or D2 receptors (**Fig. 2**). A graphical representation of the experimental paradigm is shown **Fig. 2A**, where DA was applied through iontophoresis with one of three different perfusion solutions (ACSF, ACSF+SCH23309, ACSF+Sulpiride). Our negative control (ACSF) conditions had low rates of chemoattraction (22%; **Fig. 2E,F**). The D1R antagonist SCH23390 (1 µM) conditions had 0% chemoattraction (**Fig. 2B,E,F, Supplementary Video 5**). The D2R antagonist sulpiride (600 nM) condition had no effect on DA chemoattraction compared to the ACSF control condition, with ∼75% of experiments showing no DA chemoattraction (**Fig. 2C, Supplementary Video 6**) and ∼25% showing DA chemoattraction (**Fig. 2D, Supplementary Video 7**). (**Fig. 2E-F**). Two-way ANOVA revealed a significant effect of DA receptor antagonism on chemoattraction (**Fig. 2E**; two-way ANOVA; Perfusion solution, *F*_(2, 674)_ = 19.99, *p* < 0.0001; time, *F*_(20, 674)_ = 0.0507, *p* = 1.0; interaction, *F*_(40, 674)_ = 0.0733, *p* = 1.0; DA animals n=14, slices n=22; Sulpiride animals n=6, slices n=8; SCH animals n=6, slices n=6). Together, these data suggest that NAc microglia DA chemoattraction may act through DA D1R activation and that DA D2Rs may mildly attenuate motility or not affect motility altogether.

**Figure 2:**
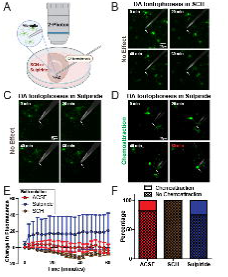
D1 and D2 receptor antagonism affects DA chemoattraction for microglia. **(A)** Schematic of experimental conditions. **(B)** Example experiment of DA iontophoresis with SCH perfusion at time points 0, 20, 40, and 60 minutes. **(C)** Example experiment of DA iontophoresis with sulpiride perfusion where no chemoattraction was observed at time points 0, 20, 40, and 60 minutes. **(D)** Example experiment of DA iontophoresis with sulpiride perfusion where chemoattraction was observed at time points 0, 20, 40, and 80 minutes. **(E)** Antagonism of D1 receptors with SCH decreases likelihood of DA chemoattraction; meanwhile, antagonism of D2 receptors with sulpiride does affect DA chemoattraction. **(F)** Antagonism of D2 receptors with sulpiride increases the likelihood of DA iontophoresis being a chemoattractant to microglia compared to SCH application.

### LPS alters microglia morphology in a similar time course to changes in DA/ATP release

The previous experiments suggest that microglia in *ex vivo* preparations are able to detect and respond to ATP and DA chemoattraction and may be responsive to DA neuron activity in the NAc through purinergic and dopaminergic receptors. We next wanted to determine whether inflammatory challenges alter DA neuron activity in addition to affecting microglia. Lipopolysaccharide (LPS), a toll-like receptor 4 agonist, is a well-established pro-inflammatory agent capable of inducing microglia to enter a reactive state characterized by functional and morphological changes including a decrease in ramification which correlates with upregulation of inflammatory markers (Qin et al., 2005). We examined the effects of LPS (1 µg/mL) on evoked DA and ATP release with FSCV across a 4-hour time course in NAc slices (**Fig. 3A-D**). Over the 4 hours, LPS increased DA release compared to ACSF control conditions (**Fig. 3C**; two-way ANOVA; Perfusion solution, *F*_(1, 378)_ = 4.682, *p* = 0.0422; time, *F*_(18, 378)_ = 4.778, *p* < 0.0001; interaction, *F*_(18, 378)_ = 3.214, *p* < 0.0001; ACSF animals n=7, slices n=9; LPS animals n=10, slices n=14), with no effect on DA clearance (Data not shown; two-way ANOVA; Perfusion solution, *F*_(1, 378)_ = 0.014, *p* = 0.9068; time, *F*_(18, 378)_ = 8.893, *p* < 0.0001; interaction, *F*_(18, 378)_ = 0.6785, *p* = 0.8328; ACSF animals n=7, slices n=9; LPS animals n=10, slices n=14). ATP release increased over the same time course, though increases were to a slightly greater extent (at 4 hrs; ATP:269.8% vs DA:210.1%) than DA (**Fig. 3A,B,D**; two-way ANOVA; Perfusion solution, *F*_(1, 324)_ = 7.653, *p* = 0.0127; time, *F*_(18, 324)_ = 3.717, *p* < 0.0001; interaction, *F*_(18, 324)_ = 2.766, *p* =0.0002; ACSF animals n=7, slices n=9; LPS animals n=9, slices n=11), with ATP clearance unaffected (Data not shown; two-way ANOVA; Perfusion solution, *F*_(1, 324)_ = 0.022, *p* = 0.8837; time, *F*_(18, 324)_ = 6.563, *p* < 0.0001; interaction, *F*_(18, 324)_ = 0.8008, *p* = 0.6995; ACSF animals n=7, slices n=9; LPS animals n=9, slices n=11). To verify that *ex vivo* NAc microglia are responsive to LPS across a similar time course, LPS (1 µg/mL) was applied to NAc core slices, and microglia were imaged using 2-photon microscopy (**Fig. 3**). LPS caused changes in microglial morphology to a more reactive state overa 4-hour time period (**Fig. 3E-G**), with decreased number of branches (**Fig. 3F**; Paired t-test; *t*_(5)_ = 3.735, *p* = 0.0135; animals n=4, cells n=6) and branch length (**Fig. 3G**; Paired t-test; *t*_(5)_ = 3.410, *p* = 0.0190; animals n=4, cells n=6). These data verify that associative *ex vivo* NAc experiments are sensitive to immune activation and indicate a similar time-course for microglia morphology changes and effects on NAc DA and ATP release.

**Figure 3:**
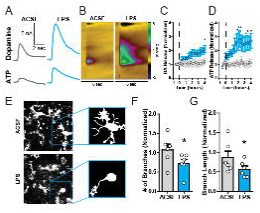
LPS induces changes in DA/ATP release in a similar time course to microglia activation. **(A)** Representative voltammetry traces of DA and ATP release before and after 4-hour LPS application with color plots shown in **(B)**. **(C)** LPS application increases evoked DA release compared to ACSF treated controls on a similar time course of microglial activation. **(D)** LPS application increases evoked ATP release compared to ACSF treated controls on a similar time course of microglial activation. **(E)** Representative LPS experiment on microglial morphology (top, ACSF baseline pre-treatment; bottom, post 4-hour LPS application; insets show representative microglia after analysis). **(F)** Four-hour LPS treatment decreases microglia normalized number of branches compared to ACSF pre-treatment. **(G)** Four-hour LPS treatment decreases microglial normalized branch length compared to ACSF pre-treatment. Asterisks *,** indicate significance levels p < 0.05 and p < 0.01, respectively, compared to ACSF.

### Glucose oxidase increases ROS and decreases DA, but not ATP release

ROS are potent immune activators (Alshehri, 2020; Mittal et al., 2014) though an examination of ROS effects on NAc microglia has not been reported. Furthermore, prior work for ROS effects on DA terminals is limited. For instance, drug-induced DA release alterations from nicotine and methamphetamine are sensitive to the ROS scavenger TEMPOL (Hedges et al., 2018). Also, NAc DA release is inhibited by supraphysiological concentrations (1.5 mM) of H_2_O_2_ (Patel & Rice, 2012), but lower, physiologically relevant ROS concentrations have not been reported. To refine upon prior work and assess effects with lower concentrations of ROS, the enzyme glucose oxidase was bath applied (**Fig. 4**) as an H_2_O_2_ generator to produce ROS from free β-D-glucose (**Fig. 4A**) (Bauer et al., 2022). Amplex Red was used to measure H_2_O_2_ levels in ACSF from increasing glucose oxidase concentrations. As anticipated, glucose oxidase increased ROS production in a concentration-dependent manner that peaked around 10 mU/mL, with H_2_O_2_ increases in the 0.5-1.5 µM range (**Fig. 4B**; one-way ANOVA; *F*_(4,14)_ = 41.98, *p* < 0.0001). Next, the same glucose oxidase concentrations were applied to brain slices while measuring evoked DA and ATP release, resulting in concentration-dependent decreases in DA release (**Fig. 4C-E**; one-way ANOVA; *F*_(4,24)_ = 3.142, *p* = 0.0437; animals n=4, slices n=5), which had no clear effect on DA clearance (Data not shown; one-way ANOVA; *F*_(4,27)_ = 0.3190, *p* = 0.8623; animals n=4, slices n=5). Glucose oxidase had mixed effects on ATP release, with increased variability and no consistent significant effect on ATP release (**Fig. 4C-D,F**; one-way ANOVA; *F*_(4,24)_ = 0.4560, *p* = 0.7668; animals n=3, slices n=5). No effect of glucose oxidase was observed on ATP clearance (Data not shown; one-way ANOVA; *F*_(4,19)_ = 0.1513, *p* = 0.9588; animals n=3, slices n=5). Next, glucose oxidase (10 mU/mL) effects on DA and ATP release were tested using a single concentration. This was done to reduce potential drift from long-duration cumulative concentration experiments that could explain the variability in ATP response in **Fig. 4F**. Consistent with our concentration response curve, significant decreases in DA release were observed with 10 mU/mL glucose oxidase (**Fig. 4G-H**; Paired t-test; *t*_(10)_ = 3.404, *p* = 0.0067; animals n=9, slices n=11). ATP release experiments still exhibited increased variability in responses after glucose oxidase, with no overall significant effect of glucose oxidase (**Fig. 4G,I**; Paired t-test; *t*_(8)_ = 0.2593, *p* = 0.8019; animals n=6, slices n=9). Thus, glucose oxidase produces H_2_O_2_ in ACSF to inhibit NAc DA release, but has mixed effects on NAc ATP release, and a single concentration application of glucose oxidase is sufficient for this DA effect.

**Figure 4:**
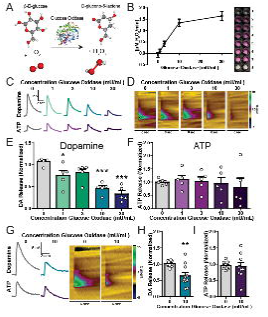
Glucose oxidase increases ROS and decreases DA, but not ATP, release. **(A)** Glucose oxidase oxidizes β-D-glucose to D-glucono--lactone to form hydrogen peroxide. **(B)** Glucose oxidase increases H_2_O_2_ production with ACSF in a dose-dependent manner. Inset: example Amplex Red experiment; 1: ACSF, no glucose oxidase (negative control), 2-4: glucose oxidase (various concentrations) in ACSF, 5-7: H2O2 (various concentrations) in reaction buffer, 8: reaction buffer, no H_2_O_2_ (negative control). **(C)** DA and ATP example traces from glucose oxidase dose response experiments. DA, but not ATP, is decreased with glucose oxidase in a dose-dependent manner. **(D)** Representative color plots from glucose oxidase dose response experiments. **(E)** Glucose oxidase decreases DA release in a dose-dependent manner. **(F)** Glucose oxidase has a varied and no clear effect on ATP release at the tested concentrations. **(G)** Representative traces of DA and ATP release before and after a single concentration (10 mU/mL) of glucose oxidase application with color plots. **(H)** 10 mU/mL glucose oxidase decreases DA release compared to ACSF pre-treatment. **(I)** 10 mU/mL glucose oxidase has no consistent effect on ATP release compared to ACSF pre-treatment but increases variability. Asterisks *, **, *** indicate significance levels p < 0.05, p < 0.01, and p < 0.001, respectively, compared to ACSF pre-treatment.

### Increases in ROS cause microglial morphology changes and slow ATP chemoattraction

IHC was performed to examine expression of ROS-sensitive TRPM2 receptors in NAc microglia (**Fig. 5A**), which was consistent with microglia in other brain regions (Jeong et al., 2017; Malko et al., 2019; Miller & Zhang, 2011) and suggests sensitivity to local ROS activity. Next, in live brain slices, 2-photon microscopy experiments were performed to measure microglial activation after application of glucose oxidase (10 mU/mL; **Fig. 5B**). This concentration was chosen based on the previous experiments that showed significant reduction in DA release at 10 mU/mL. After 1 hour, microglia had a lower number of branches compared to pre-glucose oxidase (**Fig. 5B-C**; Paired t-test; *t*_(19)_ = 2.534, *p* = 0.0202; animals n=6, cells n=20) but no change in branch length (**Fig. 5B,D**; Paired t-test; *t*_(19)_ = 0.6975, *p* = 0.4940; animals n=6, cells n=20). Thus, ROS production results in microglia morphology changes, likely through TRPM2 activation.

**Figure 5:**
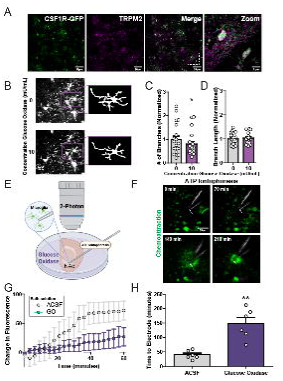
Increases in ROS cause microglial activation and slow ATP chemoattraction. **(A)** Immunohistochemical panel showing that TRPM2 receptors are located on microglia in the NAc core. **(B)** Representative 10 mU/mL glucose oxidase experiments on microglial morphology (top, ACSF baseline pre-treatment; bottom, post-glucose oxidase application; insets show representative microglia after analysis). **(C)** 10 mU/mL glucose oxidase decreases microglial normalized number of branches compared to ACSF pre-treatment. **(D)** 10 mU/mL glucose oxidase does not affect microglial normalized branch length compared to ACSF pre-treatment. **(E)** Schematic of experimental conditions for **F–H**. **(F)** Example experiment of ATP iontophoresis with glucose oxidase (10 mU/mL) perfusion at time points 0, 70, 140, and 210 minutes. **(G)** Glucose oxidase application blunts microglia chemoattraction to ATP during the first 60 minutes. **(H)** Glucose oxidase increases the time for microglia processes to reach the electrode tip. Asterisks *,** indicate significance levels p < 0.05 and p < 0.01, respectively, compared to ACSF pre-treatment.

To determine if ROS also induce changes in NAc microglia surveillance and detection of neuronal activity through ATP, glucose oxidase (10 mU/mL) was applied to live slices while examining microglia ATP chemoattraction (**Fig. 5E-F, Supplementary Video 8**). During the first hour of image acquisition, glucose oxidase application blunted ATP chemoattraction over time relative to ACSF control (**Fig. 5G**; two-way ANOVA; perfusion solution, F_(1, 10)_ = 1.812, *p* = 0.2080; time, F_(20, 200)_ = 14.11, *p* < 0.0001; interaction, F_(20, 200)_ = 4.932, *p* < 0.0001; ACSF animals/slices n=6; glucose oxidase animals/slices n=6). Consistent with this, simple linear regression showed that fluorescence increased more steeply over time in ACSF (Data not shown; slope = 1.57 ± 0.22; F_(1,124)_ = 51.78, *p* < 0.0001) than in glucose oxidase (slope = 0.45 ± 0.13; F_(1,124)_ = 12.42, *p* = 0.0006), indicating a marked slowing of microglial motility rate when ROS levels were elevated. Direct comparison of the two regressions confirmed that the slopes differed significantly (F_(1,248)_ = 19.71, *p* < 0.0001). For glucose oxidase experiments, the total recording time was increased to 3.5 hours, and complete ATP chemoattraction was eventually observed (**Fig. 5F**). The time for microglia projections to reach the electrode tip for peak fluorescence was significantly greater for glucose-oxidase-treated slices (**Fig. 5H**; Welch’s t-test; t_(5.775)_ = 4.917, *p* = 0.0030; ACSF animals/slices n=6; glucose oxidase animals/slices n=6). This suggests that in addition to altering microglia morphology, ROS generated by glucose oxidase also slow microglial motility toward ATP.

## Discussion

The present study explores the relationship between NAc DA and ATP release and local microglia activity in physiological and inflammatory conditions. NAc microglia express ATP and DA receptors and exhibit strong chemoattraction to exogenous ATP but mixed responses to DA (∼22% are sensitive to chemoattraction), suggesting the existence of microglia subpopulations with varying DA chemotactic sensitivity. Chemoattraction to DA may occur through D1-like receptor stimulation. LPS robustly modifies NAc microglia morphology, and NAc terminals release greater amounts of DA and ATP in response to LPS on a similar time-course to microglia morphology changes. Additionally, ROS change microglia morphology similar to LPS but inhibit, rather than enhance, DA release and have mixed effects on ATP release, suggesting divergent immune mechanisms for regulating microglia activity and DA terminal activity.

As resident macrophages, microglia are responsible for responding to and removing pathogens. This process is typically accompanied by a shift from ramified to more amoeboid morphology. A prime example of this is observed with exposure to the TLR4 agonist LPS, which causes microglia to reduce process length, translocate toward the pathogen, and shift from surveillant to phagocytic mode (Hines et al., 2013; Qin et al., 2005; Quan et al., 1999) while also increasing gene transcription associated with proinflammatory states to promote release of NO, H_2_O_2_, and other proinflammatory cytokines (Qin et al., 2005). Nearby cellular damage and release of DAMPs can initiate similar activity (Davalos et al., 2005; Lu et al., 2018). The present results provide evidence for complex regulation of NAc microglia behavior by local neurotransmitters including the diffuse transmitter H_2_O_2_ (Avshalumov et al., 2008). In cases where NAc DA and ATP release increase, we would expect increased microglia motility toward these terminals. In contrast, when DA and ATP release are decreased, these chemoattractants are not available and other transmitters may be prioritized.

### DA effects on microglia chemotactic motility

VTA to NAc dopaminergic projections have been extensively examined in the context of reinforcement and reward learning (Ikemoto, 2007) and have only recently been described as co-releasing ATP via VNUT-mediated vesicular packaging (Borgus et al., 2021; Ho et al., 2015). ATP is generally regarded as a DAMP chemoattractant for macrophages and microglia, and ATP effects on NAc microglia have not been described. However, ATP release is not solely a DAMP and can occur through action-potential-dependent vesicular release mechanisms (Borgus et al., 2021; Ho et al., 2015; Linderman et al., 2026) and through membrane transporters (e.g. pannexins) in response to mitochondrial stress, apoptosis, or necrosis (Chekeni et al., 2010; Vénéreau et al., 2015; Yamaguchi et al., 2014; Zhang et al., 2013). ATP on macrophages operates as a ‘find-me’ signal (Butler et al., 2021; Chiozzi et al., 2019; Haynes et al., 2006). ATP is rapidly hydrolyzed to adenosine diphosphate (ADP) and then to adenosine by ectonucleotidases in the extracellular space (Kepp et al., 2017), increasing the complexity for potential receptor targets. Indeed, the P2Y12 G_i/o_ coupled receptor, an ATP/ADP-sensitive receptor, is the most implicated for microglia chemoattraction, as a P2Y12 knockout results in potent dysregulation of chemotactic behavior in hippocampal neonate slices (Haynes et al., 2006). Prior work has also shown that ATP chemotaxis can involve a combination of G_q_ and G_i/o_ GPCR effector activity via P2Y1 and P2T_AC_ receptors (Honda et al., 2001). The P2Y6 is G_q_ coupled and is activated by UDP over ATP. A role for chemoattraction has not been established with P2Y6 and it is more implicated in phagocytosis (Plonka et al., 2019). The ionotropic P2X4 has also been implicated in microglia motility through calcium conduction and downstream cytoskeletal effectors (Ferrini et al., 2009; Nguyen et al., 2020; Verkhratsky & Burnstock, 2012). The likelihood of multiple receptor subtypes being involved in ATP chemoattraction seems ideal for triggering an immune response for selective pruning and/or clearance of damaged tissue because multiple receptors would allow for more precise detection of ATP and its byproducts.

Based on prior work indicating G_i/o_ involvement in GPCR-mediated chemotaxis (Honda et al., 2001), we originally thought that DA-induced chemotaxis would occur through the canonically G_i/o_-coupled D2-like receptors. Thus, we were surprised when DA chemotactic activity was blocked with the D1R antagonist SCH23390. Other work has shown that activation of G_s_-coupled β2 adrenergic (Gyoneva et al., 2013) and adenosine A2A receptors (Orr et al., 2009) results in process retraction via cAMP formation (Bernier et al., 2019). The finding of D1 receptors possibly mediating microglia chemoattraction highlights the complexity of receptor signaling in microglia. This unexpected finding could potentially be explained by alternate effector coupling. For example, D1-like receptors have been reported to couple to G_i/o_ effectors in some neurons (Marowsky et al., 2005), which may be a function of the cell’s expression of specific G proteins (Mönnich et al., 2024). Further, D1 and D2 heterodimers can recruit G_q_-coupled effectors (George & O’Dowd, 2007). Since DA chemotactic motility was only observed in a small subpopulation of microglia without taking into account additional controls to validate D1 receptor involvement, future studies exploring receptor effector coupling in NAc microglia may provide clues as to this novel form of chemoattraction and its role in disease and non-disease states.

### LPS effects on DA and ATP release

LPS is found on the cell walls of gram-negative bacteria and directly activates TLR4 on microglia to influence cell morphology and function (Hines et al., 2013; Lehnardt et al., 2003; Wollmer et al., 2001). LPS does not usually cross the blood-brain barrier during an infection because of its size, but it can indirectly activate microglia when administered *in vivo* through peripheral immune activity and downstream cytokine effects (Buttini et al., 1996). In the present work, LPS applied directly on brain slices caused morphology changes in microglia indicative of a pro-inflammatory, reactive state within a 4-hour period. Similarly, LPS application increased DA release across a 4-hour period. Changes in ATP release appeared sooner, with significant increases in ATP release observed in the first 1-2 hours. LPS has been shown to influence neurons, astrocytes, macrophage-lineage cells and tissue integrity, which may be how application increases DA and ATP release (Wollmer et al., 2001; Xingi et al., 2023). While the mechanisms for LPS-induced increases in DA release were not explored, it is noteworthy that the time course for effects on release matched microglia morphology changes. Based on this, one possible mechanism is that activated microglia could increase DA release through release of various cytokines. For instance, we have shown previously that the cytokines interleukin-10 (IL-10) (Ronström et al., 2023) and granulocyte colony-stimulating factor (GCSF) (Kutlu et al., 2018) both have profound, rapid excitatory effects on NAc DA release. The finding that LPS effects on ATP release were more robust than for DA, showing greater magnitude increases at earlier time points, may reflect sources of ATP beyond vesicular release from DA neurons, such as from other neurons and astrocytes (Chekeni et al., Wu et al., 2022) and from microglia themselves via lysosomal exocytosis upon TLR4 activation (Dou et al., 2012). The effects of LPS on microglia ATP chemoattraction were not explored herein, although prior work has shown that LPS can reverse the effects of ATP on microglia from acting as a chemoattractant to a repellant causing microglial process retraction rather than extension, and that this process retraction is dependent on ATP being hydrolyzed to adenosine and acting on A2A receptors on microglia (Orr et al., 2009; Gyoneva et al., 2009).

### Effect of ROS on microglia and DA release

While the source of ROS was not presently explored, striatal medium spiny neurons have been implicated in producing ROS under heavy stimulation conditions (Avshalumov et al., 2008). However, microglia are also very metabolically active and release ROS in response to pathogens (Claude et al., 2013; Herzog et al., 2019), which could be occurring in our LPS experiments, though we would predict decreases in DA release after LPS application if this were the case. Microglia also may detect and clear ROS with TRPM2, GPX, and selenoproteins (Hirrlinger et al., 2000; Jeong et al., 2017; Malko et al., 2019; Meng et al., 2019; Miller & Zhang, 2011). Indeed, we have previously reported that selenoproteins directly modulate DA transmission (Torres et al., 2021). Prior work has demonstrated that evoked DA release is reduced by high concentrations of H_2_O_2_ via changes in K_ATP_ channels (Patel & Rice, 2012). Both ROS and glucose oxidase may act on neurons, astrocytes, macrophage-lineage cells or affect tissue integrity to decrease evoked DA release. Combined with our results of increased variability in ATP release with ROS formation, activation of K_ATP_ channels by co-released ATP may contribute to some of the variability we observe in ATP responses after ROS generation, though this will need to be tested more explicitly.

DA terminals are sensitive to inhibition and activation through auto- and hetero-receptors including cholinergic, GABA, and glutamate receptors (Expósito et al., 1999; Nolan et al., 2020; Wu et al., 2000; Yorgason et al., 2017; Zhou et al., 2001) and it is possible that ROS influence activity of these local inputs to further influence DA release. Additionally, midbrain DA neurons (particularly those originating in the substantia nigra compacta) are highly sensitive to oxidative stress-induced damage (Guo et al., 2018), and the presently observed ROS effects on DA release may reflect damage to DA terminals. Thus, future work will inform on the reversibility of ROS effects on DA release. We have reported that increases in ROS underlie some of the reinforcing effects of cocaine (Jang et al., 2015) and methamphetamine (Hedges et al., 2018; Jang et al., 2017). ROS reduce DA release directly through reductions in vesicular monoamine transporter (VMAT)-mediated packaging (Hedges et al., 2018). Since ATP release is not consistently reduced by ROS, VNUT-mediated vesicular packaging appears less susceptible to ROS. The increased variability in ATP release after ROS generation suggests that in experiments where ATP is reduced by ROS, ATP packaging is influenced by DA packaging. The reduced microglia ramification observed after glucose oxidase application, coupled with our studies on ATP and DA chemoattraction, may be due to a combination of ROS effects on release of these two transmitters. We would expect a reduction in ATP and DA release to result in reduced chemoattraction and ramification, which could be a possible explanation for how glucose oxidase slowed microglia chemoattraction compared to ACSF controls. However, considering the clear expression of TRPM2 on NAc microglia, we also suspect that ROS have direct effects on microglia activity.

### Limitations

Our work provides an important step toward understanding the interplay between microglia and neurons in physiological and pathological states and provides evidence that neurotransmitters, inflammatory agents, and ROS are involved in regulating microglial and DA neuron activity. However, several important limitations of this work should be considered. First, we used murine *ex vivo* brain slice models throughout this study, which may not fully reflect how these signaling mechanisms occur *in vivo* or in the human brain. Second, as noted previously, the Csf1r-EGFP transgene mice model used does not exclusively label microglia, but also all monocyte-lineage cells including infiltrating and border-associated (e.g., perivascular and meningeal) macrophages. Csf1r is also expressed, albeit in lower levels, in certain neuronal subpopulations in the forebrain, hippocampus, and cerebellum (Hu et al., 2021), although neuronal expression in the striatum has not been demonstrated. However, a prior study did identify all Csf1r-EGFP expressing brain cells as IBA1+, supporting that this marker is a strong microglia indicator. Third, although morphological changes in microglia have been shown previously to be associated with a functional shift toward a reactive state characterized by changes in immune-related gene expression (Qin et al., 2005), morphological change alone does not necessarily signify a reactive state per se. Also, the finding that dopamine chemoattraction is observed in a subset of NAc microglia is compelling; however, further work is needed to understand the precise mechanism of this effect, including ruling out indirect mechanisms of chemotaxis such as from other signaling molecules (including potential purinergic), and ruling out possible effects of iontophoresis (e.g. effective local DA concentration, current/vehicle effects, etc). We note that, in most instances, chemoattraction was not observed in baseline images prior to iontophoretic application of ATP or DA, suggesting that tissue injury alone is not the major driver of this effect. Finally, while associations were made between DA terminal activity and microglial states in conditions of LPS- or ROS-induced inflammation, a direct causal relationship between microglia and DA and ATP activity is a subject of further research.

### Conclusions

The present work investigated the relationship between microglia and DA terminals by measuring protein expression, DA/ATP release, and microglial activation. NAc microglia express D1 and D2 receptors for DA, and TRPM2 receptors for ROS, suggesting a complex interplay within these cells for reacting to local circuit activity under normal and pathological states. ATP consistently causes microglial chemoattraction while DA appears to be selectively used by a subset of microglia. DA bath application does not impair ATP-induced chemoattraction, and we observed little evidence for DA chemorepulsion. Application of LPS increases DA and ATP release and alters microglia morphology across a similar time course. Glucose oxidase induces dose-dependent increases in ROS and causes reductions in DA releasebut mixed effects on ATP release. ROS also altered microglia morphology and slowed process motility to ATP. The present study highlights a major gap in our understanding and complexity in neuroimmune interactions between microglia and DA terminals and is relevant for understanding the role of microglia in conditions of DA dysfunction such as attention, mood and substance use disorders.

## Declarations

### Ethical Approval and Consent to Participate

Not applicable.

### Consent for Publication

Not applicable.

### Availability of Data and Materials

All data needed to evaluate the conclusions herein are present in the research article and/or Supplementary Materials.

### Author Contributions

**HAW**: Conceptualization, Formal Analysis, Investigation, Writing – Original Draft, Visualization, Supervision, Project Administration. **JAW**: Writing – Review and Editing, Visualization, Formal Analysis. **EBT**: Investigation, Supervision. **ERW**: Investigation, Supervision. **LHF**: Formal Analysis, Investigation, Visualization, Supervision. **LRH**: Formal Analysis, Investigation, Visualization, Supervision. **BJP**: Investigation, Visualization. **STJ**: Software, Investigation. **SCL**: Investigation, Visualization. **CJG**: Investigation, Supervision. **DDL**: Investigation. **CAS**: Conceptualization, Methodology, Resources. **JMH**: Methodology, Resources. **SCS**: Conceptualization, Writing - Review & Editing. **JTY**: Supervision, Funding Acquisition, Resources, Conceptualization, Methodology, Writing - Review & Editing.

## Acknowledgements

Special thanks to Rachel H. Webb for feedback and editorial assistance.

