## Supplementary Figure 1 for "Reactive oxygen species alter ex vivo dopaminergic and purinergic signaling and microglia physiology in the nucleus accumbens"

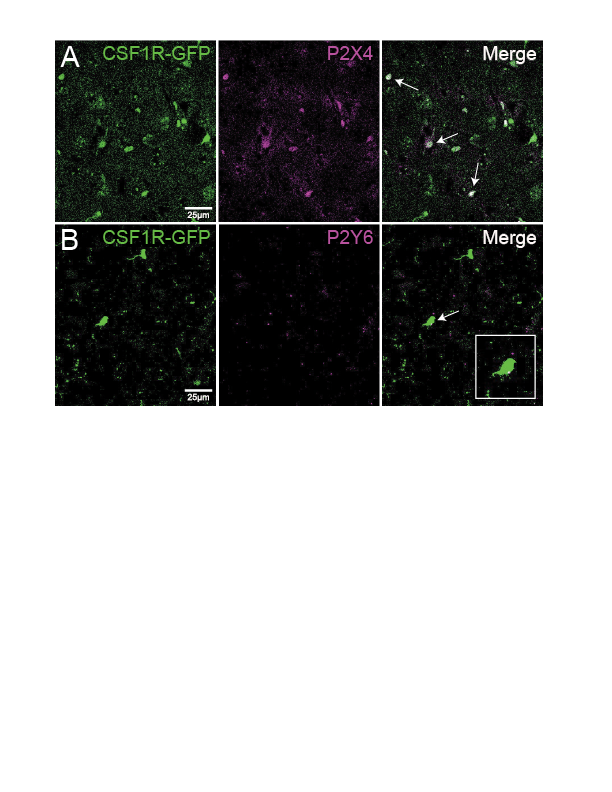

***Supplementary Figure 1****: Purinergic receptor expression in NAc microglia.* **(A)** Immunohistochemical panel showing ATP P2X4 receptor labeling on NAc microglia. Left panel: CSF1R-GFP+ microglia, middle panel: P2X4 receptor labeling, right panel: merge of P2X4 and GFP labeling. Arrows indicate microglia with clear P2X4 labeling. **(B)** Immunohistochemical panel showing UDP sensitive P2Y6 receptor labeling on NAc microglia. Left panel: CSF1R-GFP+ microglia, middle panel: P2Y6 receptor labeling, right panel: merge of P2Y6 and GFP labeling and an inset zoom of one microglia with P2Y6 labeling. P2Y6 labeling is present but reduced compared to P2X4 labeling.
